# UDP-glucosyltransferase 71C4 regulates seed development by redistributing phenylpropanoid metabolism in cotton

**DOI:** 10.1101/2023.11.09.566436

**Authors:** Yiwen Cao, Zegang Han, Lu He, Chujun Huang, Jinwen Chen, Fan Dai, Lisha Xuan, Sunyi Yan, Zhanfeng Si, Yan Hu, Tianzhen Zhang

**Author notes:** Corresponding author: Tianzhen Zhang.

## Abstract

Seeds are important for plant reproduction, and hence it is important to identify genes regulating seed development. Here, we focused on a member of the glycosyltransferase (GT) family, UDP-glucosyltransferase 71C4 (*UGT71C4*), which influences the seed width, seed length, and yield traits (lint percentage and seed index) by regulating the lignin and flavonoid pathways of phenylpropane metabolism, affects the oil and protein contents of mature seeds, and controls the seed size by regulating cell proliferation. Overexpression of *UGT71C4* leads to seed enlargement by activating expression of peroxiredoxins and the flavonoid metabolism pathway; induces accumulation of ROS, which promotes cell proliferation; and significantly improves yield traits, with the seed index increasing from 10.66 to 11.91 and protein increasing by 5.34%, but oil content decreasing 8.98%. Conversely, knockout of *UGT71C4* causes seeds to become smaller; the lignin metabolism pathway to be activated, especially enzymes relating to lignin synthesis leading to increased ectopic deposition of lignin in the ovule and constrained ovule growth and development; and significant improvement of yield traits, with lint percentage increasing from 39.62% to 41.94%, seed index decreasing from 10.66 to 8.60, protein content decreasing 4.28%, and no significant change in oil content. Our research provides new insights into seed size regulation through UDP-glucosyltransferase, providing potential methods for improving plant yield.

## INTRODUCTION

Seeds are the unique reproductive bodies of gymnosperms and angiosperms, and as such playing a causal role in the continuation of a species and serving serve as the foundation of agricultural production. Most plant seeds bear the heavy responsibility of species reproduction and diffusion, among which with crop seeds also provide providing food and necessary nutrients for humans and livestock. Mature seeds are rich in various nutrients, including carbohydrates, storage proteins, and lipid storage compounds; ultimately, providing seeds provide approximately 70% of the energy intake for organisms on Earth. For instance, soybean protein is the world’s main source of plant protein and an abundant source of oil as well (Li et al., 2019), while rice is the source of carbohydrates for half of the world’s population (Song et al., 2007).

The diversity and complexity of plant secondary metabolites is a consequence of many post-translational modifications such as methylation, glycosylation, hydroxylation, and acylation (De Bruyn et al., 2015; Tiwari et al., 2016; Louveau and Osbourn, 2019; Ye et al., 2022; Cardenas et al., 2019). Glycosylation is the last modification to be made on plant products, and as such, glycosyltransferases (GTs) are central to plant growth and metabolism, moreover, glycosylation is a core modification in regulating biological activities, which has led to the development of glycodiversity. Notably, the cell walls of plant seeds contain abundant carbohydrate polymers, and changes in sugar metabolism abundance can alter the accumulation of substances during seed development (Gomez et al., 2009). A plant secondary product glycosyltransferase consensus sequence (called the PSPG box) is the critical binding domain recognized by UDP-glucosyltransferase, which has the ability to transfer glucuronic acid from uridine diphosphate (UDP)-glucose as the sugar donor to proteins, lipids, and small-molecule secondary metabolites, thereby influencing their stability, solubility, and bioactivity(Le Roy et al., 2016; Mackenzie et al., 1997). The PSPG box consists of about 40 amino acids and is located near the C-terminal sequence of UGTs; within the box, two short sequences WAPQV and HCGWNS are highly conserved, being present in 95% of GTs. Notably, GTs lose their activation ability if the conserved histidine and glutamic acid are mutated (Jones, 2000). *Arabidopsis* UGTs can be divided into 115 families (Coutinho et al., 2003; Gachon et al., 2005), of which GT1 comprises 107 glycosyltransferases and is further divided into 16 subgroups (groups A-P). Members of a subgroup share greater than 60% similarity and have similar substrates and donors (Brazier-Hicks et al., 2018). With the evolution of higher plants, the E group has become the largest GT subgroup; it includes *UGT71*, *UGT72*, and *UGT88*, among many others. E group members *UGT71B6* and *UGT71C5* are able to recognize and glycosylate abscisic acid (ABA) and ABA-related metabolites (Priest et al., 2006; Liu et al., 2015). Overexpression of *UGT72E1*, *UGT71E2*, and *UGT71E3* accelerates the accumulation of coniferyl alcohol 4-O-glucoside in phenylpropanoid metabolism (Lanot et al., 2006). Additionally, *OsGSA1* encodes a UDP-glucosyltransferase that regulates grain size by regulating cell proliferation and expansion (Dong et al., 2020).

Overall, more than 100,000 secondary metabolites have been found across plant species in past decades (Gachon et al., 2005). These metabolites evolved over a long period of time and play important roles in cellular damage, stress tolerance, and protection from insect invasion (Goodman et al., 2004; Hu et al., 2013; Iven et al., 2012). Plant secondary metabolites are extensively utilized as signal molecules, for example to mediate auxin movement and plant development (Peer and Murphy, 2007; Hu et al., 2021; Mateo-Bonmati et al., 2021). Multiple metabolic pathways, including lipid (Yang et al., 2017; Niu et al., 2009), sugar (Wang et al., 2021a; Gomez et al., 2009) and phenylpropanoid (Besseau et al., 2007), are involved in seed formation and development. Phenylpropane metabolism in particular is one of the main sources of secondary metabolites, including as it does the flavonoid and lignin pathways. Flavonoids are involved in many aspects of plant growth and development, such as in pathogen resistance, UV damage repair, pollen growth, and seed coat development; meanwhile, lignin is the second most abundant polymer on Earth, accounting for around 30% of the organic carbon in the biosphere.

Lignin is a composite phenolic polymer deposited in the secondary cell walls of all vascular plants. Xylem cell differentiation is a key step in the formation of secondary cell walls, serving to enhance the mechanical strength of xylem conduits and provide hydrophobicity (Wilkerson et al., 2014). Lignin monomer biosynthesis involves a series of hydroxylation, methylation, and other modification reactions to form basic lignin complexes. Natural lignin polymers are formed from p-coumaryl, coniferyl, and sinapyl alcohol, which respectively produce p-hydroxyphenyl (H), guaiacyl (G), and syringyl (S) units (Ralph et al., 1997; Voxeur et al., 2015). Spatial and temporal control of the lignification process is extremely important; lignification is a metabolically costly process that requires a large carbon skeleton, and plants do not have mechanisms for degrading lignin. Therefore, plants must balance the need for lignification with the resources available for synthesizing lignin polymers. In addition, because lignin limits cell wall expansion, lignification must follow cell division and expansive growth. Lignin is also implicated in seed development. Negative regulation of lignin synthesis and deposition in rice promotes the formation of larger grains (Jiang et al., 2019). The protein SH5 is involved in the lignin synthesis pathway and inhibits lignin deposition during early anther development to promote anther development (Yoon et al., 2014). The *Arabidopsis* mutant *dgd1* exhibits decreased photosynthesis, altered chloroplast morphology, and lignification in the cap cells of the phloem. Further analysis of hormone and lipidomic revealed a significant decrease in digalactosyldiacylglycerol content, leading to the occurrence of lignification (Lin et al., 2016). To date, transcription factors that stimulate lignin synthesis have mostly been identified among the MYB and NAC families. *PtMYB4* and *PtMYB8* from *Pinus taeda* participated in the phenylpropanoid metabolism pathway and are activators of secondary cell wall deposition in conifers (Bomal et al., 2008). Overexpression of *AtMYB46*, an Arabidopsis homologue of *PtMYB4* and *PtMYB8*, has been shown to activate expression of genes involved in the biosynthesis of xylan, cellulose, and lignin, and hence participates in the network regulating Arabidopsis secondary wall synthesis (Zhong et al., 2007). NAC family proteins encoded by *NST1*, *NST2*, and *NST3* directly target *AtMYB46* and its homologs, which are redundant regulators of stem and anther secondary wall synthesis in *Arabidopsis*. *MtNST1* is the only NST gene to have been identified in the model legume *Medicago truncatula*, and Mtnst1 mutants exhibit secondary wall synthesis defects in the stem and anthers (Zhao et al., 2010).

Seed development in general has been well-studied, but the mechanism of action of glycosyltransferase in seed development is still unclear. Here we report a gene encoding UDP-glucosyltransferase (*UGT71C4*) to be involved in cotton seed development. Overexpression or knockout of *UGT71C4* leads to changes in the flavonoid and lignin content of seeds due to altered expression of genes in the phenylpropane metabolism and lignin synthesis pathways, which ultimately impacts seed size, lint percentage, and protein and oil content. These findings could lay a solid foundation for further seed formation and development research in plants and could have benefits for crop yield improvement.

## Results

### Isolation and characterization of *UGT71C4*, a gene predominantly expressed during ovule development

Our RNA-seq results (Hu et al., 2019) identified that UDP-glucosyl transferase 71C4 (*UGT71C4*, *GH_D05G0690*), a gene of length 1458 bp that encodes 486 amino acids, was predominantly expressed in ovule tissues at 10, 15, and 20 days postanthesis (DPA), suggesting it plays roles in ovule development and cotton seed maturation (**Fig. 1A**). To gain insights into the function of *UGT71C4*, we performed a phylogenetic analysis against the flavonoid UGTs of *Arabidopsis* and found that *UGT71C4* was largely resembled the UGT71 group, which are known as flavonoid catalytic enzymes (**Supplemental Figure S1 and Table S1**). A conserved region named the PSPG box, which is widely associated with the synthesis of plant secondary metabolites and binding of UDP sugar donors, was detected in the C-terminus amino acid sequence of *UGT71C4* (**Fig. 1B and Fig. 1C**). The protein spatial structure of *UGT71C4* contains a pair of tightly linked and face-to-face β/α/β Rossmann-like domains, termed a GT-B fold (**Fig. 1D**) which structure is responsible for recognizing and binding donors and acceptors through the resulting loose cleft (Lairson et al., 2008). All in all, *UGT71C4* from Glycosyltransferase family 1 possesses a GT-B fold, is predominantly expressed during ovule development, and has a typical catalytic domain PSPG box; it therefore might be responsible for catalyzing terpenes, flavonoids, and anthocyanins, as is the case for other GT-B UGTs.

**Figure 1.**
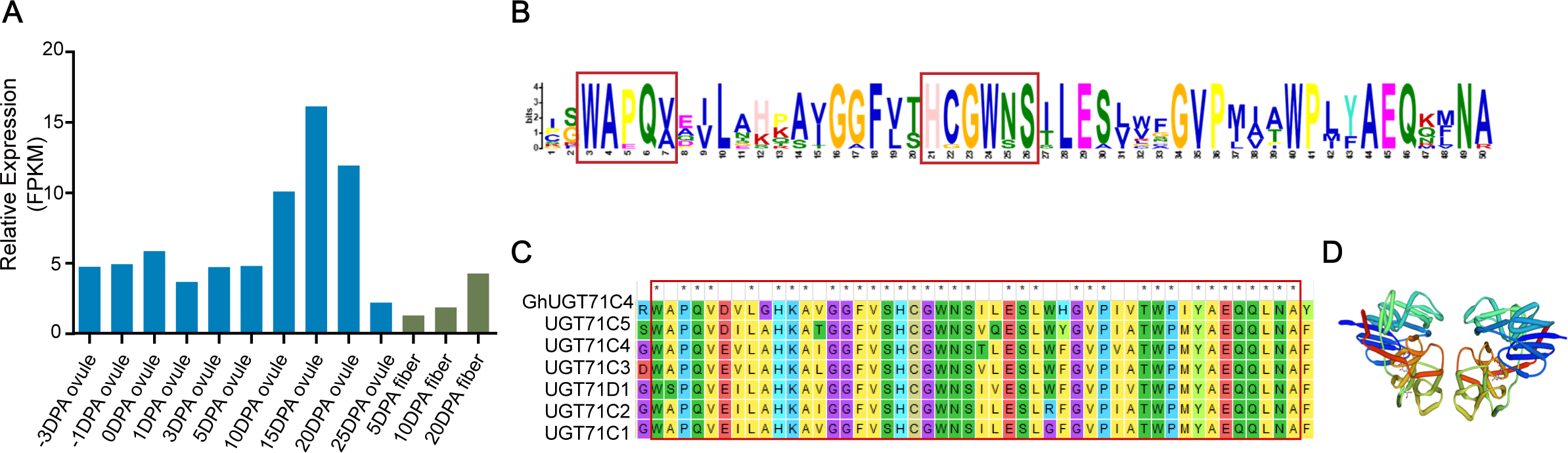
Phylogenetic tree, protein structure and characterization of *UGT7lC4*. **A**) *UGT7lC4* transcript level (FPKM) from -3 to 35 DPA in ovule and at 5, 10, and 20 DPA in fiber. **B)** Protein sequences of the conserved plant secondary product glycosyltransferase (PSPG) motif in all UGTs, which contains a conserved domain of 44 amino acids and two short highly conserved sequences indicated by a red box. **C)** Multiple alignment of cotton *UGT7lC4* and *Arabidopsis* UGTs that are its near neighbors in the phylogenetic tree The conserved domain of the PSPG box is indicated by a red box, and * represents conserved amino acids. **D)** GT-B enzyme structure showing the two BlalB Rossmann-like domains, which are oriented face-to-face and are not very closely related. The active site is located in the gap between them.

### *UGT71C4* pleiotropically regulates seed and lint development, and also oil and protein content

To gain detailed insights into the function of *UGT71C4* in cotton, we employed clustered regularly interspaced short palindromic repeats (CRISPR)/CRISPR-associated nuclease 9 (Cas9)-mediated target mutagenesis to produce five different knockout lines (*UGT71C4-KO*). The single-guide RNAs (sgRNAs) consisted of 20 bp sequences, and mutations were induced in a ∼20 bp region upstream of a protospacer adjacent motif (PAM) in the cotton receptor line W0. After producing T1 homozygous lines, two homozygous and CRISPR-associated nuclease 9 protein free transgene lines, named *UGT71C4-KO-3* and *UGT71C4-KO-5* and harboring a 1-bp thymine (T) deletion and a 1-bp adenine (A) insert respectively, were used for further experiments (**Fig. 2A, Supplemental Figure S2A**). Both lines exhibited a reduction in seed size (**Fig. 2B and 2C**); seed widths were significantly narrower (*UGT71C4-KO-3* 4.57±0.98 mm and *UGT71C4-KO-5* 4.36±1.02 mm vs. W0 4.86±0.90 mm) (**Fig. 2D**), and mature seed lengths were dramatically shorter (*UGT71C4-KO-3* 8.13±1.42 mm and *UGT71C4-KO-5* 8.57±0.64 mm vs. W0 10.34±0.51 mm) (**Fig. 2E**). As was expected, *UGT71C4-KO-3* and *UGT71C4-KO-5* also exhibited seed index reduction, by 19.30% and 15.97% respectively, as compared with W0 (**Fig. 2F**). However, lint percentage (LP) was significantly higher (5.8%) in *UGT71C4-KO-5* than in the receptor W0 (**Fig. 2G**).

**Figure 2.**
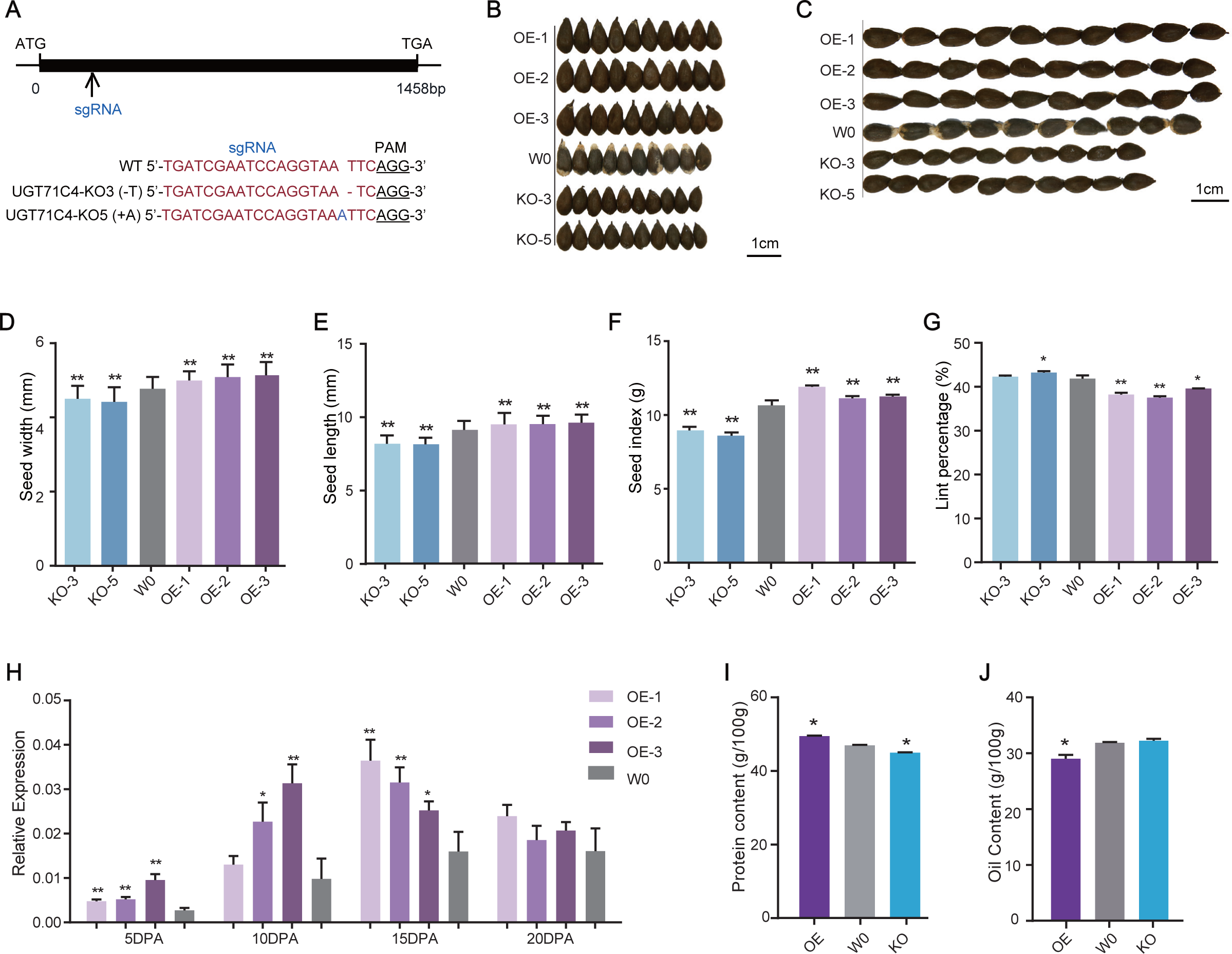
*UGT7lC4* affects seed size and weight. **A)** CRISPRlcas9 mediated mutagenesis of *UGT7lC4*. Top: Schematic diagram of *UGT7lC4*; the CRISPRlCas9 target site is indicated by an arrow. Exons and introns are represented by black boxes and black lines, respectively. Bottom: Alignment of receptor W0 and mutant sequences containing the target site. In the receptor sequence, the target sequence adjacent to the underlined PAM is indicated in red. The newly generated *UGT7lC4-K0-3* and *UGT7lC4-K0-5* mutants harbored a 1-bp T-deletion and a 1-bp A-insertion, respectively. **B)**, **D)** Seed width and **C)**, **E)** seed length in transgenic plants compared with receptor plants. *UGT7lC4-0E-l*, *UGT7lC4-0E-2*, and *UGT7lC4-0E-3* are overexpression lines; *UGT7lC4-K0-3* and *UGT7lC4-K0-5* are knockout lines; W0 is the receptor. Scale bar represents 1 cm. **F)** Seed index of the various *UGT7lC4* lines. **G)** Lint percentage of the various *UGT7lC4* lines. **H)** Relative expression of *UGT7lC4* in 5, 10, 15, and 20 DPA ovules. **I)** Mature seed protein content of *UGT7lC4-0E*, *UGT7lC4-K0*, and W0. **J)** Mature seed oil content of *UGT7lC4-0E, UGT7lC4-K0*, and W0. Values in Figure2 represent means ± s.d. *P < 0.05 and **P < 0.01 indicate significant difference by two-tailed Student’s t-test (seed width and length, n = 50; others, n=3).

Next, the full-length coding sequence of *UGT71C4* was cloned into a vector with the CaMV 35S promoter and used to generate overexpression lines. More than 11 independent overexpression lines were identified among the T1 homozygotes, of which the top three lines with highest expression were further investigated; these are referred to as *UGT71C4-OE-1*, *UGT71C4-OE-2*, and *UGT71C4-OE-3* (**Supplemental Figure S2B**). Expression during ovule development was quantified by qRT-PCR, which showed the overexpression lines to have significantly increased *UGT71C4* abundance in 5, 10, 15, and 20 DPA ovules compared with W0 (**Fig. 2H**). Seeds of the overexpression lines were significantly (*P*<0.01) wider and longer (*UGT71C4-OE-1* w: 5.02±0.78 mm, l: 10.93±0.63 mm; *UGT71C4-OE-2* w: 5.20±0.92 mm, l: 10.69±0.50 mm; and *UGT71C4-OE-3* w: 5.22±0.83 mm, l: 9.81±0.93 mm vs. W0 w: 4.86±0.90 mm, l: 10.34±0.51 mm) **(Fig. 2D and Fig. 2E**). Larger grain and seed size contributes to higher crop yield, to a certain extent. The overexpression line *UGT71C4-OE-1* demonstrated the most significant increase of seed index, at 11.78% more than W0; it was followed by *UGT71C4-OE-3* at 5.56% and then *UGT71C4-OE-2* at 4.47% **(Fig. 2F)**. However, LP was decreased by 2.22%, 2.17%, and 4.64% in *UGT71C4-OE-1*, *UGT71C4-OE-2*, and *UGT71C4-OE-3*, respectively (**Fig. 2G**). There was no stable and significant change of fiber length, elongation, strength or macronaire value in either the overexpression or knockout lines (**Supplemental Table S2**).

Finally, we selected the overexpression and knockout lines with the most obvious phenotypes (*UGT71C4-OE-1* and *UGT71C4-KO-5* respectively; henceforth *UGT71C4-OE* and *UGT71C4-KO*) and measured their protein and oil contents. Compared to the receptor, the protein content in *UGT71C4-OE* seeds was significantly increased by 5.34%, going from 45.83 g/100 g to 48.27 g/100 g, while oil content was significantly reduced by 8.98%, from 31.89 g/100 g to 29.03 g/100 g. Conversely, the protein content of mature seeds of the *UGT71C4-KO* line was significantly decreased by 4.28% and the oil content increased by only 1.16%, which did no achieve significance (**Fig. 2I and Fig. 2J**). In summary, *UGT71C4* pleiotripically regulates seed and lint development along with oil and protein content, without affecting fiber quality.

### *UGT71C4* regulates seed development by controling cell proliferation

To understand the effects of *UGT71C4* excess and depletion during seed development, we tracked changes in ovule development from 5 DPA to 20 DPA. There was no significant difference among the plants at 5 DPA or 10 DPA, but great differences in seed size appeared at 15 DPA to 20 DPA (**Fig. 3A**). Additionally, scanning electron microscopy (SEM) of the outer epidermal cells of ovules at -1 DPA revealed a certain degree of correlation between epidermal cell number and ovule size. Compared with W0, there were fewer epidermal cells observed in *UGT71C4-OE*, and more in *UGT71C4-KO* (**Fig. 3B and Fig. 3C**). *UGT71C4-OE* also produced a greater number of ovule inner seed coat cells compared to W0, resulting in cells of smaller size that were tightly arranged with small cell gaps (**Fig. 3D and Fig. 3E**). In terms of endoplasmic cell cross section, the individual cells of *UGT71C4-KO* were larger in area and more loosely arranged.

**Figure 3.**
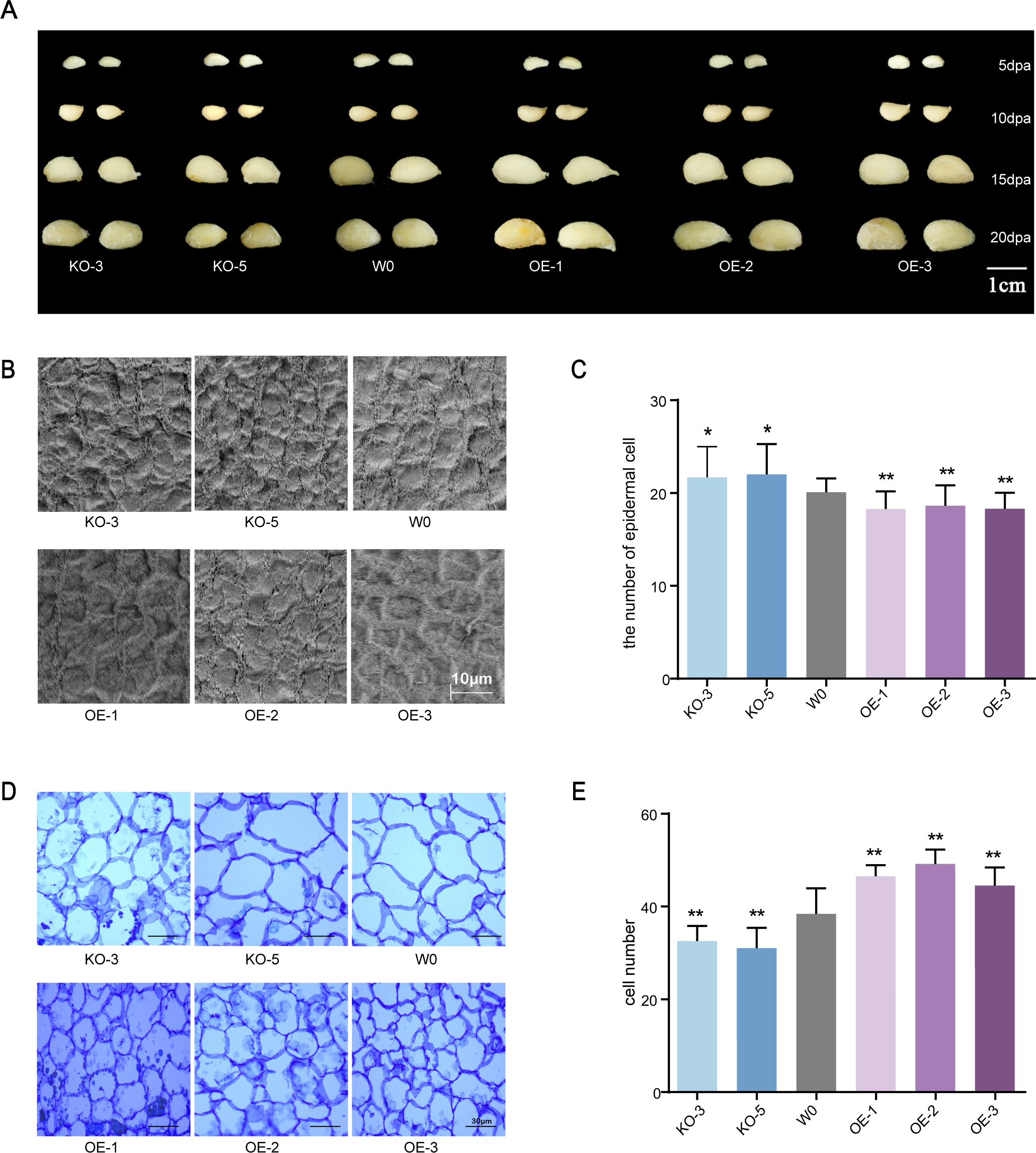
*UGT7lC4* controls seed size by regulating seed cell proliferation. **A)** Ovule size in transgenic lines from 5 to 20 DPA. Scale bar represents 1 cm. **B)** Scanning electron micrographs of ovule outer epidermal cells at -1 DPA. Scale bar represents 10 µm. **C)** Number of outer epidermal cells (n = 20-25) at 10 DPA. Scale bar represents 30 µm. **D)** Images of parenchymal cells of the ovule inner integument from *UGT7lC4-0E-l*, *UGT7lC4-0E-2*, *UGT7lC4-0E-3*, *UGT7lC4-K0-3*, *UGT7lC4-K0-5* and W0. **E)** Number of inner integument cells in the same area (n=6-10).

Ultimately, the *UGT71C4-OE* line possessed more inner seed coat cells and larger seeds, whereas *UGT71C4-KO* had smaller seeds with fewer inner seed coat cells that were sparsely arranged. Therefore, we inferred that *UGT71C4* is involved in seed development by regulating cell proliferation and expansion.

### Metabolomics revealed *UGT71C4* activates the flavonoid metabolism pathway to regulate seed development

Through UPLC-MS/MS detection and database analysis, we detected a total of 97 differential metabolites in 15 DPA ovules of *UGT71C4-OE* compared to W0, of which 60 metabolites were down-regulated, including ferulic acid, sinapic acid, cordycepin, and isoscopoletin, and 37 metabolites were up-regulated, including naringenin, dihydrokaempferol, dihydroquercetin, and cinnamic acid. In *UGT71C4-KO*, a total of 112 differential metabolites were detected, of which 23 metabolites were down-regulated, including dicaffeoylquinic acid-O-glucoside, and formononetin-7-O-glycoside, and 89 metabolites were up-regulated, such as eriodictyol, p-coumaryl alcohol, and phenyl acetate (**Supplemental Table S3**). KEGG pathway enrichment analysis of the differential metabolites identified enrichment of “Phenylpropanoid biosynthesis”, “Flavonoid biosynthesis”, “Flavone and flavonol biosynthesis”, and “Biosynthesis of secondary metabolites” pathways (**Supplemental Figure S5**). The phenylpropane metabolic pathway produces a large number of secondary metabolites, mainly flavonoids and monolignols. As compared with W0 and *UGT71C4-KO*, the 15 DPA ovules of *UGT71C4-OE* had apparently higher flavonoid levels (**Fig. 4A**), excepting the flavonoid glycoside which was accumulated in ovules of the overexpression line.

**Figure 4.**
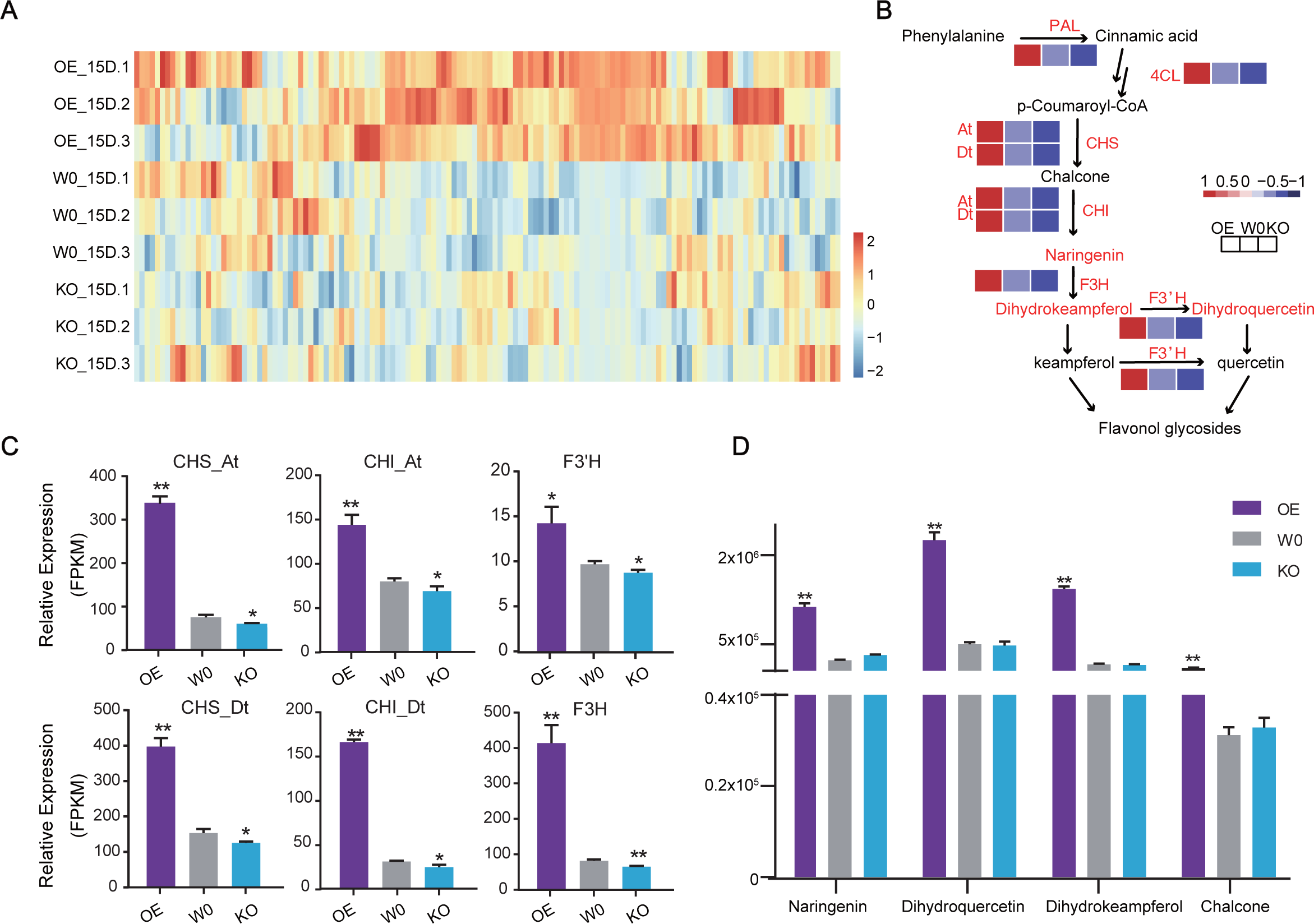
Flavonoid metabolism was changed in seeds of *UGT7lC4-0E* transgenic plants. **A)** Heatmap of the relative flavonoid and lignin contents in ovules at 15 DPA for *0E*, *K0*, and W0 (n=3 biological replicates) based on widely targeted metabolomic data. **B)** Mapping of flavonoid metabolism pathways to visualize overall differences in 15 DPA ovules of *0E*, *K0*, and W0 plants. Relative expression of key catalytic enzymes is shown in the box; text in red indicates significant difference compared to W0, and text in blue no significant difference. **C)** Relative expression levels of flavonoid-biosynthesis-related genes. **D)** Differential metabolites in the flavonoid metabolic pathway (n = 3 biological replicates). P < 0.05 and P < 0.01 indicate significant difference by two-tailed Student’s t-test.

Phenylalanine ammonia lyase (PAL) acts as an entry enzyme in the phenylpropane metabolic pathway to introduce more reduced carbon into basal phenylpropane metabolism, while 4-coumarate-CoA ligase (4CL) is an enzyme of the flavonoid and lignin branches, and hence is involved in the synthesis of flavonoids and cell-wall-bound phenolics. Both *PAL* and *4CL* showed significantly higher expression in *UGT71C4-OE*, but significantly lower expression in *UGT71C4-KO* **(Fig. 4B)**. RNA-seq data validated that the four major genes involved in flavonoid synthesis, chalcone isomerase (CHI: *GH_A13G0216*, *GH_D04G0140*), chalcone synthase (CHS: *GH_A09G0002*, *GH_A02G0270*, *GH_D02G0295*), flavanone 3-hydroxylase (F3H: *GH_A12G0596*), and flavonoid 3′-monooxygenase (F3’H: *GH_A12G2012*), are likewise significantly increased in *UGT71C4-OE* but significantly decreased in *UGT71C4-KO* (**Fig. 4C**). Surprisingly, the changes in metabolite levels correspond clearly with changes in the expression of these genes for both *UGT71C4-OE* and *UGT71C4-KO*. In particular, chalcone content was significantly increased by 3.29 fold in 15 DPA ovules of *UGT71C4-OE*, but not detected in ovules of *UGT71C4-KO*. Naringenin, the downstream product of chalcone, also exhibited 4.89 fold greater accumulation in ovules of the overexpression line. Similar trends were observed for dihydroquercetin and dihydrokaempferol, downstream products of the flavonoid pathway, which were significantly accumulated by up to 4.53 and 8.95 fold respectively in overexpressing ovules as compared with the receptor W0 (**Fig. 4D**).

Taken together, the above results support that *UGT71C4* actively participates in the flavonoid metabolism branch of the phenylpropanoid metabolism pathway. In 15 DPA ovules of *UGT71C4*-*OE* plants, flavonoid metabolism and synthesis enzymes are more active; moreover, relative to plants with loss of *UGT71C4* function, those overexpressing *UGT71C4* prefer to flux phenylpropane metabolism products into the flavonoid pathway over the lignin pathway.

### *UGT71C4* involves in the lignin synthesis, cell proliferation and ROS content

To further elucidate the molecular mechanism by which *UGT71C4* influences seed development, we conducted transcriptome sequencing (RNA-seq) and widely targeted metabolomic assays in 15 DPA ovules of *UGT71C4-OE* and *UGT71C4-KO* and their receptor W0. RNA-seq analysis identified a total of 4359 differentially expressed genes (DEGs) in *UGT71C4-OE* compared with receptor W0, including 2200 up-regulated genes and 2159 down-regulated genes; meanwhile, *UGT71C4-KO* exhibited only 279 DEGs, with 72 up-regulated and 207 down-regulated (**Supplemental Figure S3 and S4**). Gene Ontology (GO) enrichment analysis identified terms such as “cell wall”, “cell wall organization”, “cell wall biogenesis”, and “peroxidase activity” to be enriched among DEGs in both the overexpression and knockout lines (**Fig. 5A and Fig. 5B**), suggesting that *UGT71C4* regulates cell wall structure and peroxidase activity. GO analysis showed that a large number of genes differentially expressed in *UGT7C4-OE* and *UGT71C4-KO* were enriched in terms related to “cell wall” and “cell wall organization”, as mentioned above (**Fig. 5A and Fig. 5B**). Lignin is one of the main components of the cell wall, and significant accumulation of lignin was observed in mature seeds of *UGT71C4-KO,* but no significant difference in *UGT71C4-OE* (**Fig. 5C**) as compared to the receptor W0. Moreover, four typical lignin synthesis activators, *MYB4*, *MYB83*, *MYB46_A*, and *MYB46_D*, were predominantly expressed in *UGT71C4-KO*, while the lignin synthesis inhibitor *MYB8* was predominantly expressed in *UGT71C4-OE*. In addition to members of the MYB family, the NAC family gene *NST1*, a lignin synthesis activator, showed decreased transcription level in the ovules of *UGT71C4-OE*, while lignin-synthesis-related genes were significantly active in *UGT71C4-KO* (**Fig. 5D**). Finally, lignin monomer synthesis occurs primarily through the lignin metabolism pathway in phenylpropane metabolism, and transcriptional levels of catalytic enzymes in this pathway were also changed with perturbation of *UGT71C4* expression. Especially, hydroxycinnamoyl-CoA:shikimate hydroxycinnamoyl transferase (HCT: *GH_D05G1187*) and caffeic acid O-methyltransferase (COMT: *GH_A12G2656*) were both significantly decreased in *UGT71C4-OE*, but significantly increased in *UGT71C4-KO* (**Fig. 5E**).

**Figure 5.**
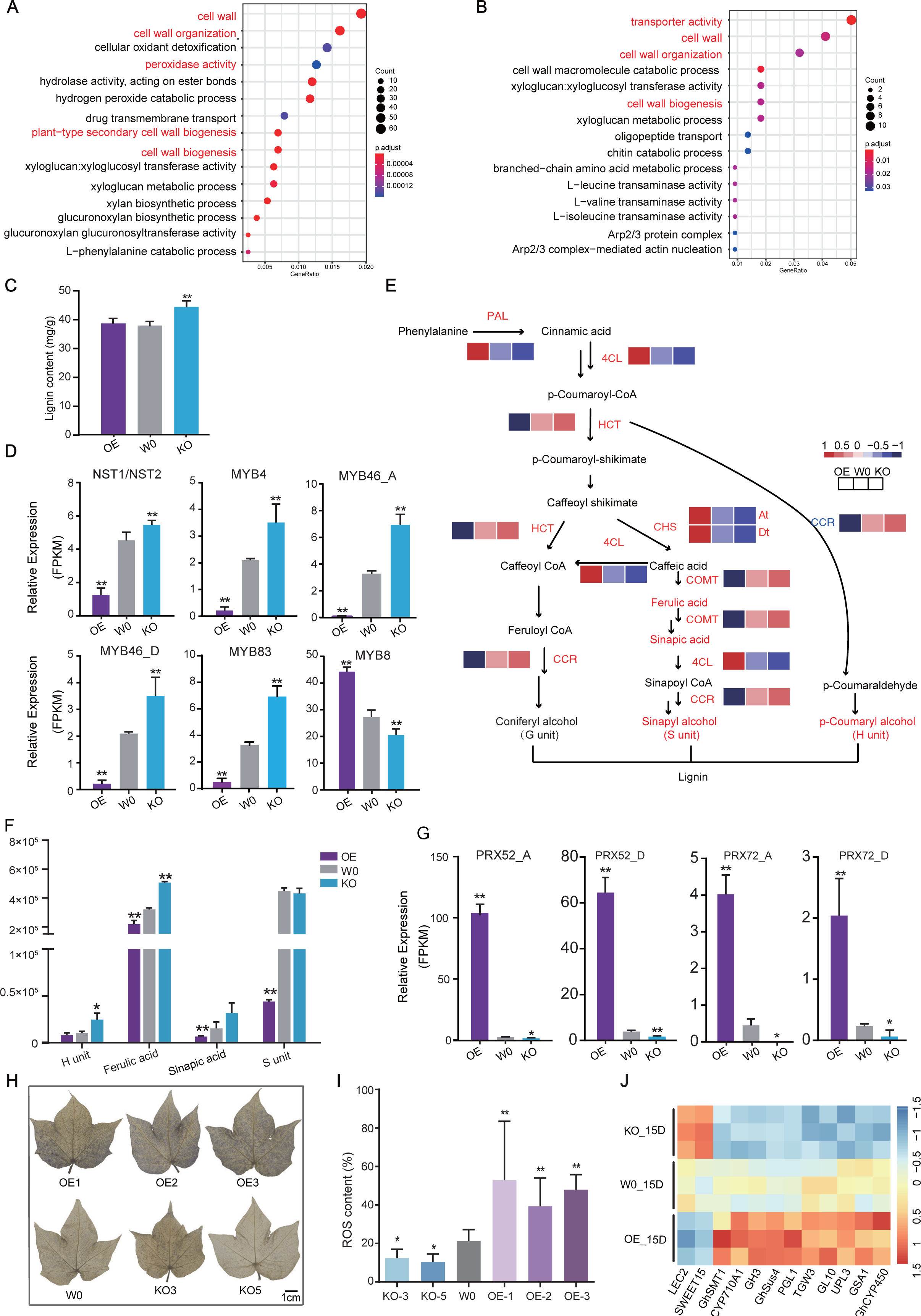
Altered expression of lignin metabolism, ROS content, and cell proliferation associated genes in *UGT7lC4*. **A), B)** GO enrichment analysis of genes differentially expressed genes of in *UGT7lC4-0E* (A) and *UGT7lC4-K0* (B) **C)** Mature seed lignin content in *UGT7lC4-0E*, *UGT7lC4-K0*, and W0 (n = 3 biological replicates). P < 0.05 and P < 0.01 indicate significant difference by two-tailed Student’s t-test. **D)** Relative expression levels of lignin-biosynthesis-related genes. **E)** Mapping of lignin metabolism pathways to visualize overall differences in 15 DPA ovules of *UGT7lC4-0E, UGT7lC4-K0*, and W0 plants. The relative expression of key catalytic enzymes is shown in the box; text in red represents significant difference compared to W0, and text in blue no significant difference. **F)** Differential metabolites in the lignin metabolic pathway. **G)** Relative expression levels of *PRX72_A*, *PRX72_D* and P*RX52_A, PRX52_D* in *UGT7lC4-0E*, *UGT7lC4-K0*, and W0. **H)** and **I)** Comparison of oxygen radical production in leaves of *UGT7lC4-0E-l, UGT7lC4-0E-2*, *UGT7lC4-0E-3,* W0, *UGT7lC4-K0-3*, and *UGT7lC4-K0-5* using NBT staining (n=3 biological replicates). **J)** Regulation of the relative expression of genes involved in cell division. P < 0.05 and P < 0.01 indicate significant difference by two-tailed Student’s t-test.

Considering changes in lignin content and lignin and lignin monomer synthesis genes in mature seeds, the levels of the monolignol and the intermediate products were significantly increased in the *UGT71C4-KO*, but decreased in *UGT71C4-OE* ovules (**Fig. 5F**). For example, considering the main lignin monomers p-coumaryl alcohol (H unit) and sinapyl alcohol (S unit), the former had 2.40-fold higher content in *UGT71C4-KO* ovules compared to those of W0, but *UGT71C4* the overexpression reduced it to only 0.78 times the control value; meanwhile, the latter one exhibited no significant change in *UGT71C4-KO* ovules has no significant changes, while *UGT71C4-OE* ovules contained only 0.09 times as much S monomer as W0. The Similar results were observed for the precursors of sinapyl alcohol, ferulic acid and sinapic acid; was similarin particular, ferulic acid accumulated in *UGT71C4-KO* ovules was to 1.58 times higher than the level in the receptor.

RNA-seq and GO term enrichment analysis revealed the term “peroxidase activity”, which is associated with cell proliferation, to be enriched only in DEGs of *UGT71C4-OE*, and not those of *UGT71C4-KO* (**Fig. 5A and Fig. 5B**). The DEGs annotated with this term include *PRX72_A* and *PRX52_A*, which are homologous to *AtPRX72* and *AtPRX52*, sharing 88% and 79% amino acid similarity respectively (**Supplemental Figure S6A and S6B**). As illustrated in **Fig. 5G**, *PRX72_A*, *PRX72_D* and *PRX52_A*, *PRX52_D* are transcribed at a significantly higher level in *UGT71C4-OE* and a significantly lower level in *UGT71C4-KO*. Nitroblue tetrazolium (NBT), which can be oxidized to a dark blue precipitate by superoxide anions, was used to quantify oxygen radicals in the leaves of *UGT71C4-KO*, *UGT71C4-OE*, and W0. This staining revealed accumulation of a large number of superoxide anions in *UGT71C4-OE* leaves, but few in *UGT71C4-KO* leaves (**Fig. 5H and Fig. 5I**).

We additionally observed differential expression of many genes related to cell growth and development, such as *OsTGW6* (*GH_A01G2007*) has been shown to control the cellularization stage of endosperm to increase grain weight (Ishimaru et al., 2013). The gene *GmSWEET10a* and the CYP450 family (Yang et al., 2013; Tang et al., 2016; Zhao et al., 2016; Wang et al., 2015; Liu et al., 2020) had been shown to affect seed size, oil content, and protein levels in soybean (Duan et al., 2022); the gibberellic acid signaling pathway gene *GH_A05G2747* (*GL10*) (Zhan et al., 2022); sucrose synthesis pathway genes like *GH_A05G0363* (*GhSus4*) not only mainly involved in the synthesis and storage of starch and protein (Jiang et al., 2012; Ruan et al., 2003; Liu et al., 2020); the phytosterol biosynthetic gene *GH_A08G0610* (*GhSMT1*) (Suo et al., 2021); and the UDP-glucosyltransferase *GH_D05G3357* (*GSA1*), which is known to control grain size and abiotic stress tolerance (Dong et al., 2020); meanwhile, *DA1* and the homologous gene *GW2* are key genes that positively regulate cell proliferation in *Arabidopsis thaliana* and maize, but negatively regulate grain weight and grain width in rice (Xie et al., 2018; Li et al., 2008; Achary and Reddy, 2021).and other genes related to seed development had a remarkable increased or decreased (**Fig. 5J**). We selected for validation by qRT-PCR the HECT E3 ligase *UPL3*, which has proven involvement in seed development, and the transcription factor *PGL1* (*PRE1*), a member of the basic helix-loop-helix (bHLH) family that has roles in cell division and cotton plant development (Zhang et al., 2009). The results validated that the expression of these two genes is altered by *UGT71C4* perturbation, being significantly increased in *UGT71C4-OE* and decreased in *UGT71C4-KO* ovules at 5 DPA, 10 DPA, and especially 15 and 20 DPA (**Supplemental Figure S7**).

Taken together, in addition to significant lignin deposits in *UGT71C4-KO*, genes and enzymes related to lignin synthesis were more active compared to the receptor; conversely, they were not significantly altered in seeds of the overexpression line. When *UGT71C4* is overexpressed or knocked out, the balance of the phenylpropanoid metabolism pathway is broken, metabolic flow is preferentially directed towards lignin metabolism rather than the flavonoids pathway, more lignin than flavonoids is accumulated in the ovule, and ROS levels are altered, all of which resulted in alteration of ovule cell proliferation and eventually seed development.

## DISCUSSION

The gene *UGT71C4* has huge potential in improvement of cotton production. With its knockout in *UGT71C4-KO*, LP was increased from 39.62% to 41.94% and seed oil content was slightly increased by 1.16%. Meanwhile, with its overexpression in *UGT71C4-OE*, seed index was increased by 11.78%, ovule inner seed coat cells were more closely arranged and more numerous, mature seeds were longer and wider, and protein content exhibited a significant increase of 5.34% (**Fig. 2 and Fig. 3**). In our prior study of 318 cotton accessions (Hu et al., 2022), we observed a negative relationship between cottonseed oil and protein contents, which is consistent with the current findings in transgenic lines. Nonetheless, improvements in SI or LP would benefit cotton production, while improving protein or oil content promotes practical use of cottonseeds. In that light, *UGT71C4* could be a candidate gene for future studies aimed at improving yield parameters or seed composition.

### Altered flavonoid pathways and seed development

Joint analysis of RNA-seq and a widely targeted metabolomics assay revealed that the seeds of *UGT71C4-OE* accumulated more flavonoids in their ovules, including chalcone, naringenin, dihydroquercetin, and dihydrokaempferol (**Fig. 4, Supplemental Figure S4**). Some key synthases (CHI, CHS, F3H, F3’H) in the flavonoid metabolic pathway were also significantly elevated (**Fig. 4C**). The first three steps of phenylpropane metabolism underlie the entirety of the pathway, and provide precursors for flavonoid and lignin metabolism. Phenylalanine ammonia lyase (PAL), cinnamic acid 4-hydroxylase (C4H), and 4-coumarate-CoA ligase (4CL) are generally the core enzymes of phenylpropane metabolism. PAL introduces metabolic fluxes into phenylpropane metabolism (Zhang and Liu, 2015). In previous studies, mutation of chalcone synthase in *Zea mays* and *Petunia hybrida* was found to block the first step of the flavonoid pathway, resulting in growth defects in pollen tubes and hindered root hair development (Taylor and Grotewold, 2005; Buer and Muday, 2004). The flavonoid content in sweet potato (*Ipomoea batatas*) leaves is reportedly directly proportional to leaf size and the number of epithelial cells, and hence greater flavonoid content can promote leaf development (Gao et al., 2023). Thus, flavonols can directly promote cell growth and support storing more energy. *CHS* enacts the first step in catalyzing the flavonoid pathway, which results in a series of flavonoid products. Silencing of *CHS* in soybean transgenic materials has been shown to result in a lack of flavonoids, which inhibits nodulation formation (Wasson et al., 2006). In our current study, these three key enzymes were significantly affected by perturbation of *UGT71C4*. As the cotton seed is also a major site of nutrient storage, we infer that more metabolic fluxes flow into flavonoid metabolism with overexpression of *UGT71C4*. As moderate accumulation of flavonoids has been shown to contribute to cell proliferation, the accumulation of flavonoids in ovules tends to promote seed development. Our findings thus provide new insights into the roles of flavonoids.

Researchers have intensively studied the glycosylation substrates of *Arabidopsis thaliana* UGT71 family members, who are involved in the glycosylation of phenylpropane metabolic intermediates with high substrate specificity for low-molecular-weight compounds such as flavonoids. In particular, enzymes in the UGT71 family are all capable of glycosylating two hydroxycoumarins, scopoletin and esculetin (Lim, 2002; Lim et al., 2001), as previously determined *in vitro*, with *UGT71C1* showing catalytic activity towards the 3-OH of hydroxycoumarins, hydroxycinnamates, and flavonoids (Lim et al., 2003; Lim et al., 2008). Our unrooted evolutionary tree indicates that *UGT71C4* belongs to group E, which is the same as the *Arabidopsis* UGT71 family, and the sequence similarity of functioning PSPG boxes is extremely high (**Fig. 1B and Fig. 1C**). Combined with existing research results, we hypothesized that *UGT71C4* in cotton has the same ability to catalyze certain substances in the phenylpropane metabolic pathway as the *Arabidopsis* UGT71 family, and hence its genetic perturbation results in the redistribution of the phenylpropane metabolic pathway and changes of the key enzyme activities and metabolite contents observed in transgenic plants.

### Ectopic accumulation of lignin limits seed development

Research in rapeseed has shown that phenolic compounds can affect seed weight and size, as well as alter the internal structure of the seed and the overall metabolic level of the plan (Clauss et al., 2011). Following repression of the lignin biosynthetic gene *HCT* in *Arabidopsis*, the metabolic flux from lignin synthesis flows to the flavonoid accumulation pathway, resulting in a dwarf phenotype without a floral stem (Besseau et al., 2007; Hoffmann et al., 2004). In the current work, several related synthases (*HCT*, *COMT*, *CCR*) were transcribed at significantly higher levels in *UGT71C4-KO* ovules, but suppressed in *UGT71C4-OE*. The intermediates ferulic acid, sinapic acid, p-coumaryl alcohol, and sinapyl alcohol also accumulated significantly in *UGT71C4-KO* ovules (**Fig. 5E and Fig. 5F**). Accumulation of such key enzymes and metabolites in the lignin pathway may be part of the reason enlargement of *UGT71C4-KO* ovules is constrained.

Lignin biosynthesis is also regulated by MYB and NAC transcription factors. *NST1* and *NST2* are two typical factors that promote secondary wall thickening; overexpression of either leads to ectopic deposition of the secondary cell wall in normal parenchyma cells (Mitsuda et al., 2005; Zhao et al., 2010). Hyperactivation of MYB activators leads to severe transcript accumulation for genes in the lignin synthesis pathway, ectopic deposition in plant secondary walls, small curled leaves, and smaller floral organs (Patzlaff et al., 2003; Zhong et al., 2007). With further research on the MYB family, factors with inhibitory functions have also been discovered (Bomal et al., 2008). Notably, some NAC and MYB genes that promote lignin synthesis were significantly increased in *UGT71C4-KO* and significantly repressed in *UGT71C4-OE*, whereas the opposite trends were observed for a lignin synthesis repressor (**Fig. 5C and Fig. 5D**).

The spatial and temporal control of lignification is extremely important for plants, as excessive deposition of lignin in tissues undergoing active division or elongation could lead to decreased cell wall tension, which limits cell division and elongation and ultimately has adverse effects on plant growth and development (Zhao, 2016). Based on existing evidence, we inferred that the smaller seeds of *UGT71C4-KO* plants may be attributed to altered transcription of lignin-synthesis-related genes, enhanced lignin deposition, erroneous ectopic expression, and more metabolic flux in phenylpropane metabolism flowing into the lignin metabolic pathway rather than flavonoid metabolism. Ectopic deposition of lignin in the ovule led to decreased ductility of the secondary wall, which limited the expansion of endoplasmic cells, ultimately decreasing individual cell volume and reducing seed size.

### *UGT71C4* regulates seed size through ROS levels and control of cell proliferation genes in cotton

It is well established that peroxidase family genes play vital roles throughout the entire plant life cycle (Passardi et al., 2004). During leaf development, peroxidases not only down-regulate lignin content but also regulate ROS levels, thereby altering cell proliferation and expansion (Shigeto et al., 2013; Li et al., 2003; Blee et al., 2003; Lu et al., 2014; Wang et al., 2021b). However, reducing the ROS level below a certain value maight inhibit cell proliferation, affecting normal differentiation and defense (Schieber and Chandel, 2014). ROS stimulates cell growth by increasing cell extensibility through the presence of the hydroxyl (⋅OH) radical in the cell, and elongation of cotton fibers is dependent on ROS to regulate cell wall relaxation and retard the increase in cell wall rigidity (Ruan et al., 2001; Gapper and Dolan, 2006). The results in this work showed that peroxidase family genes have significantly altered expression in *UGT71C4-OE* and *UGT71C4-KO* **(Fig. 5G)**, supporting that normal levels of ROS are crucial for cell growth and development. Combining the observed patterns of lignin deposition and ROS accumulation, it is clear that ROS promotes cell development to a certain extent. Specifically, in *UGT71C4-KO*, hydroxyl (⋅OH) radicals were less active in tissues with more lignin deposition, whereas in *UGT71C4-OE* lines, hydroxyl radicals accumulate significantly in tissues with less lignin deposition, where they increase cellular extensibility and promote cell division, and particpate in coordinating cellular activities (**Fig. 5H and Fig. 5I**). Consistent with previous studies, we found *UGT71C4* to be involved in phenylpropane metabolism in addition to energy flow; furthermore, we observed it to finely regulate genes involved in grain size regulation

## Conclusion

We propose a working model in which metabolic changes in upland cotton seeds are regulated through *UGT71C4* glycosyltransferase, affecting seed size, oil content, and protein content. In *UGT71C4-OE,* the phenylpropanoid metabolism pathway is induced in seeds, flavonoid metabolism is active, and key catalytic enzymes in the flavonoids pathway such as CHS, CHI, and F3’H are significantly upregulated at the transcriptional level. Substances such as naringenin, dihydrokeampferol, and dihydroquercetin are appropriately accumulated in the ovule. In addition, genes related to cell proliferation are significantly upregulated, promoting ovule cell expansion, and the seed protein content is significantly increased (**Fig. 6**). In contrast, in *UGT71C4-KO* seeds, enzymes related to the flavonoids pathway and its products are significantly inhibited, and the lignin synthesis pathway is instead induced. Lignin monomers and intermediate products are overaccumulated in the ovule, and lignin polymerization ability is significantly improved. Ectopic deposition of lignin in the ovule reduces the ductility of the cell wall and limits seed growth and development; genes relating to secondary wall synthesis are significantly induced, resulting in the formation of smaller grains; and protein content is significantly reduced (**Fig. 6**).

**Figure 6.**
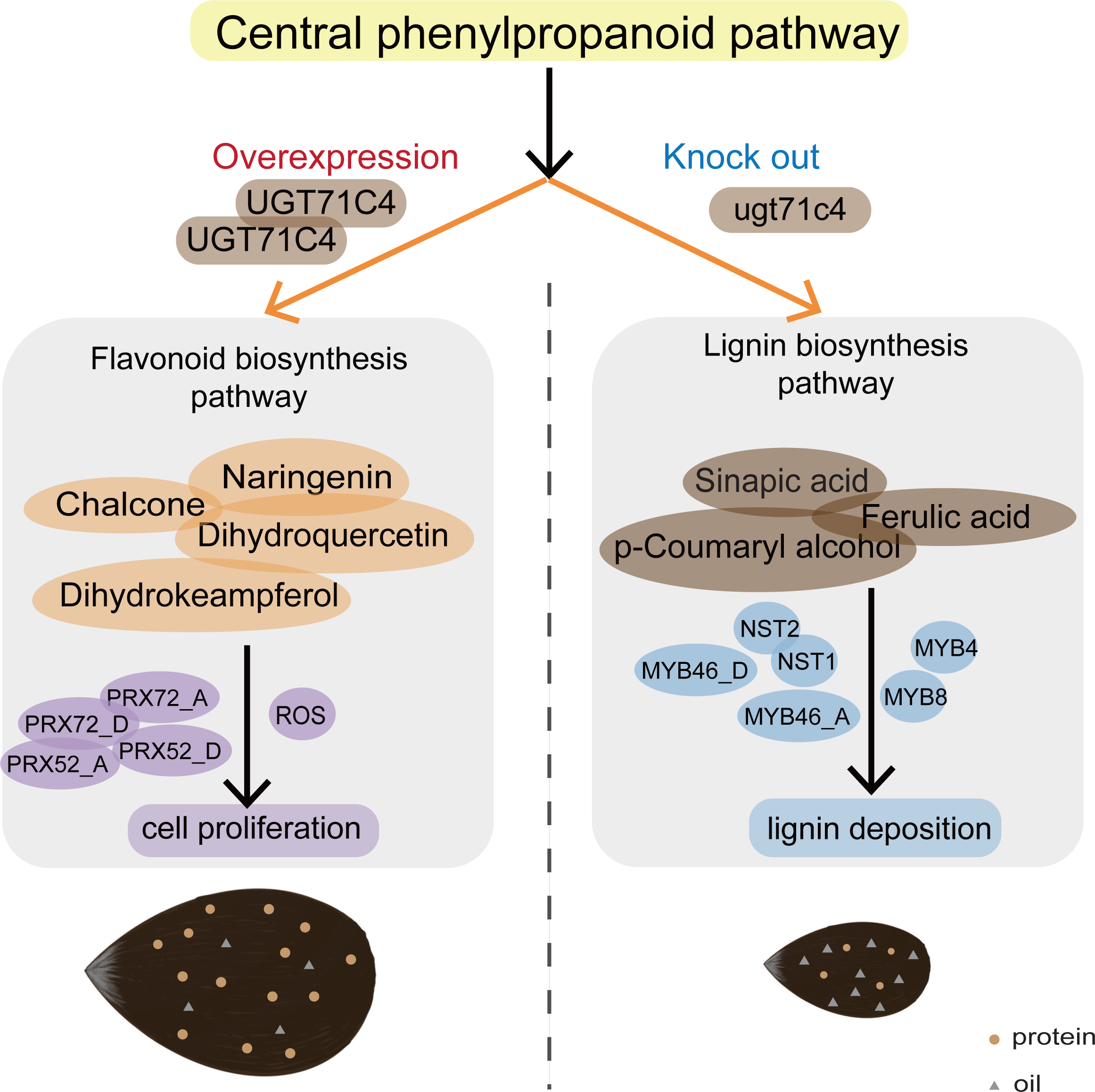
Working model of *UGT7lC4* regulation of seed size. When *UGT7lC4-0E* increases the metabolic flux from basic phenylpropane metabolism to flavonoid metabolism, the expression of core flavonoid synthesis enzymes is increased significantly; appropriate flavonoids are accumulated in ovules to promote cell growth and development; the expression of genes related to cell differentiation and proliferation is advantageously regulated; and ROS is accumulated in the plant, which further promotes cell proliferation in ovules and ultimately increases seed length and width and hence the yield. In *UGT7lC4-K0* ovules, more metabolic flux flows from basal phenylpropane metabolism to lignin metabolism, which results in greater lignin monomer and intermediate synthesis; significantly increased expression of related core enzymes; induction of genes related to lignin synthesis; and significant accumulation of lignin in the ovules, which in turn results in lignification of the cell wall, reduced cell ductility, and hence constrained seed development.

## MATERIALS AND METHODS

### Vector construction, plant genetic transformation, plant culture

The full-length coding sequence of *UGT71C4* was cloned into a vector containing the CaMV 35S promoter. The CRISPR-cas9 system was used to edit genomic material, with sgRNAs targeting *UGT71C4* designed using the website CRISPR-P 2.0 (http://crispr.hzau.edu.cn/CRISPR2/). Vectors containing cloned inserts were transformed into cotton using *Agrobacterium tumefaciens*-mediated method with the strain LBA4404 (CAT#: AC1030). Genomic DNA inserts from overexpression lines were amplified by PCR to confirm transformation, while knockout lines were confirmed by Hi-TOM (**Supplementary Table S4**). Confirmed overexpression, knockout, and receptor plants were cultivated in the fields of Hainan, Dangtu, and Hangzhou of China. Seeds from homozygous plants were collected after maturity for subsequent experiments. The seed index (weight of 100 seeds) was measured for over 300 seeds from 25 bells.

### Phylogenetic analysis and sequence alignment

UGT amino acid sequences were obtained from previous reports in *Arabidopsis thaliana* (Brazier-Hicks et al., 2018) (**Supplementary Table S1**). Multiple sequence alignment of UGTs was performed using Clustal W. MEGA X was used to construct the neighbor-joining (NJ) tree with 1000 bootstrap replicates. Multi-sequence alignment between *UGT71C4* and other known UGT proteins was performed using BioXM (http://202.195.246.60/BioXM/).

### Plant RNA extraction and quantitative reverse transcription polymerase chain reaction (qRT-PCR

Total RNA of different cotton tissues was extracted with the RNAprep pure Plant Kit (DP432), then converted into cDNA using HiScript II Q RT SuperMix for qPCR (+gDNA wiper) (Vazyme: R223-01). A LighterCycler96 (Roche) and ChamQ Universal SYBR qPCR Master Mix (Vazyme #Q711) were used to quantify gene expression by the 2^-ΔΔ^ CT method. PCR primers are given in **Supplementary Table S4**.

### Transcriptome sequencing

Transcriptome sequencing was performed by Novogene (Beijing, China). First, total RNA sequences were assessed using the Fragment Analyzer 5400 (Agilent Technologies, CA, USA). Next, the transcriptome sequencing library was prepared and sequencing libraries were generated using the NEBNext® UltraTM RNA Library Prep Kit for Illumina® (NEB, USA). All samples were clustered using the TruSeq PE Cluster Kit v3 cBot HS (Illuminia) on the cBot Cluster Generation System. The library formulation was sequenced using the Illumina Novaseq 6000 platform to generate paired-end reads of 150 bp. Raw data converted from the Illumina platform was subjected to quality control before subsequent analysis. The transcriptome analysis method was as described previously (He et al., 2022).

### Widely targeted metabolomics methods

The widely targeted metabolomics assay was performed by Metware (Wuhan, China). Flowers were labeled on the day of flowering and ovules from different lines were harvested 15 days postanthesis (DPA). The ovule samples were freeze-dried using a vacuum freeze-drying machine (Scientz-100F), then were crushed with a mixed mill (MM 400, Retsch) and zirconia beads at 30 Hz for 1.5 minutes. The resulting freeze-dried powder (100 mg) was dissolved in 1.2 mL of 70% methanol solution, incubated at room temperature for three hours with vortexing for 30 seconds every 30 minutes (a total of six times), and then placed overnight in a refrigerator at 4 ℃. Prior to UPLC-MS/MS analysis, the extract was centrifuged at 12000 rpm for ten minutes, then filtered (SCAA-104, pore size 0.22 μm; ANPEL, Shanghai, China, http://www.anpel.com.cn/). A UPLC-ESI-MS/MS system was then used to analyzed the sample extracts. Analytical conditions for UPLC were as follows. The column was an Agilent SB-C18 (1.8 µm, 2.1 mm * 100 mm). The mobile phase consisted of solvent A: pure water containing 0.1% formic acid, and solvent B: acetonitrile containing 0.1% formic acid. The gradient program had initial conditions of 95% A and 5% B, graded linearly to 5% A, 95% B within nine minutes, maintained the composition of 5% A, 95% B for one minute, graded to 95% A and 5% B within 1.1 minutes, and then maintained that composition for 2.9 minutes. The flow rate was set to 0.35 mL/min; the column oven to 40 ℃; and the injection volume to 4 μL. The wastewater was alternately connected to the ESI triple quadrupole linear ion trap (QTRAP) mass spectrometer LIT and triple quadrupole (QQQ) scans were obtained on a triple quadrupole linear ion trap mass spectrometer (Q TARP), the AB4500 Q trap UPLC/MS/MS System. This system was equipped with an ESI Turbo ion spray interface, which operates in both positive and negative ion modes and is controlled by the Analyst 1.6.3 software (AB Sciex). The operation parameters of the ESI source were: ion source, turbine spray; source temperature, 550 ℃; ion spray voltage (IS), 5500 V (positive ion mode)/-4500 V (negative ion mode); ion source gas I (GSI), gas II (GSII), and curtain gas (CUR) flows of 50, 60, and 25.0 psi, respectively; and collision activation dissociation (CAD) was high. The instrument was turned and calibrated with a 10 and 100 μmol/L polypropylene glycol solution in the QQQ and LIT modes. QQQ scanning was an MRM experiment conducted with the collision gas (nitrogen) set to medium. Further DP and CE optimization was carried out for the transition of a single MRM. Monitor a specific set of MRM transitions for each period based on the metabolites eluted during this period.

### Histochemical analysis and scanning electron microscopy

Ovules (15 DPA) were fixed overnight in 50% ethanol, 5% glacial acetic acid, and 5% formaldehyde at 4 °C, then dehydrated in an ethanol series. For histochemical analysis, after fixation with xylene, the ovules were embedded in an extracellular matrix (Sigma Aldrich) and sliced into 8-mm-thin sections using a rotary slicing machine (Leica). The prepared tissue sections were then stained with toluidine blue and observed under a Carl Zeiss light microscope. Subsequently, critical point drying (Leica) of the spikelet shell, gold sputter coating, and observation under scanning electron microscopy (Hitachi) was performed. Cell size and number were measured using ImageJ, and images were collected using the Leica application suite (V 4.2).

### Lignin content determination

Samples (0.05 g) were ground into powder, then combined with 5 mL of 1% acetic acid and mixed well. The mixture was centrifuged, and the sediment retained. Ethanol ether mixed solution (3 mL) was then added to the precipitate, mixed thoroughly, allowed to incubate for three minutes, and then centrifuged, again with the precipitate retained. After drying the precipitate, 1.2 mL of 72% sulfuric acid was added to the precipitate, mixed well, and let stand at room temperature for 16 hours (overnight) to dissolve all the cellulose. The next day, 4 mL of distilled water was added to the centrifuge tube, mixed well, and incubated in a boiling water bath for five minutes. Afterwards, 2 mL of distilled water and 0.2 mL of 10% barium chloride solution were added; the solution was shaken well, then centrifuged, retaining the precipitate. The precipitate was next washed with 5 mL distilled water, then resuspended in 4 mL of 10% sulfuric acid and 4 mL of 0.025 mol/L potassium dichromate solution. After mixing, the precipitate was heated in a boiling water bath for 15 min with shaking. After cooling, all substances in the centrifuge tube were transferred to a conical flask. Residual substances in the centrifuge tube were washed out with 15-20 mL water, which eluate was also added to the flask. For titration, 2 mL of 20% KI solution was added to the conical flask, and the solution was calibrated with sodium thiosulfate to the end point. Next, 0.4 mL of 1% starch solution was added, and titration with sodium thiosulfate was continued until the solution turned from blue to bright green. At that end point, the titration volume was recorded, the blank volume V0 was calculated using the same method, and the lignin content was determined.

## FUNDING INFORMATION

This study was financially supported by grants from the Fundamental Research Funds for the Central Universities (226-2022-00100) and the NSFC (32130075).

## SUPPLEMENTAL DATA

**Supplemental Figure S1.** Phylogenetic tree analysis of *UGT71C4*.

**Supplemental Figure S2.** The detection of transgenic cotton.

**Supplemental Figure S3.** Venn diagrams of total DEGs in *UGT71C4-OE* and *UGT71C4-KO* ovules.

**Supplemental Figure S4.** Genes differentially expressed genes of in *UGT71C4-OE* and *UGT71C4-KO* ovules.

**Supplemental Figure S5.** The KEGG enrichment pathway enrichment analysis of significant differential metabolites of in *UGT71C4* transgenic ovules.

**Supplemental Figure S6.** Amino acid sequence similarity analysis and ROS content.

**Supplemental Figure S7.** Relative expression levels of genes related to seed size development related genes in *UGT71C4* transgenic ovules.

**Supplemental Table S1.** List of the genes used in phylogenetic tree.

**Supplemental Table S2.** Statistic of *UGT71C4* transgenic lines.

**Supplemental Table S3.** Up and down regulated metabolites in *UGT71C4* transgenic lines.

**Supplemental Table S4.** List of the primers used in this study.

